# Comparative genomics reveals insights into phylogenomic taxonomy and potential algae-bacteria interactions of novel versatile *Mameliella alba* strain LZ-28 isolated from highly-toxic marine phycosphere

**DOI:** 10.1101/2021.05.17.444442

**Authors:** Wen-zhuo Zhu, Fei-fei Xu, Yun Ye, Qiao Yang, Xiao-ling Zhang

## Abstract

Phycosphere harbors cross-kingdom interactions with significant ecological relevance for harmful algal blooms (HAB) and phycotoxins biosynthesis. Previously, a new red-pigmented bacterium designated as strain LZ-28 was isolated from phycosphere microbiota of typical HAB dinoflagellate *Alexandrium catenella* LZT09 which is a vitamin B_12_ auxotroph and produces high levels of paralytic shellfish poisoning toxins (PST). Strain LZ-28 exhibited obvious growth-promoting activity toward its algal host, along with the production of active bioflocculanting exopolysaccharides (EPS). But the phylogenetic affiliation and genomic potential of this versatile bacterium has not yet been elucidated. In this study, we carried out combined taxonomic and phylogenomic analysis to clarify the taxonomic classification of strain LZ-28. The obtained 16S rRNA phylogeny revealed close taxonomic relationship between strain LZ-28 and other *Mameliella alba* members. Additional calculations of key phylogenomic parameters, average nucleotide identity (ANI), the average amino acid identity (AAI) and the digital DNA-DNA hybridization (dDDH) values based on genomes of strain LZ-28 and type strain of *Mameliella alba* were all exceeded the limit of species circumscription. Collectively considering the phenotypic and biochemical characterizations, strain LZ-28 was therefore identified as a new member of *Mameliella alba.* Furthermore, based on the genomic evidence, potential algae-bacteria interactions of strain LZ-28 with host algae LZT09 were elucidated through the associations with photosynthetic and antioxidant carotenoids, supplying of bacterial VB_12_ to auxotroph host, and versatile EPS serving for bacterial colonization and nutrient exchange during their interactions, along with stress response systems to defense oxidative stress and quorum sensing (QS) signals benefited survival for bacteria in the symbiotic system. Comparative genomics shed light on similar genomic features between *M. alba* strains, revealed potential close associations of strain LZ-28 with its algae host, and further enriched the genomic repertoire of interactions between phycosphere microbiota and algal host LZT09.

## Introduction

Phycosphere, the boundary of phytoplankton holobionts, is a unique microscopic niche for algae-bacteria interactions (ABI) [1,2]. These dynamic interplay between algae and their closely-associated phycosphere microbiota harbors cross-kingdom exchanges of diverse nutrients, secondary metabolites, infochemicals, and gene transfer agents (GTA) through their stable symbiotic relationship. The exchange for ammonium, antimicrobials, bioactive compounds (such as vitamin B_12_), detoxification of reactive oxygen species, inorganic carbon (Ci), and nitrogen fixation of bacteria may result from the readily available sources of carbon and organic matters by algal host [3]. Also, those complex inter-species and intra-species associations plays foremost roles in primary production, and biogeochemical cycles and the microbial loop in the oceans [4]. Moreover, it’s also one key player to regulate the biosynthesis of paralytic shellfish poisoning toxins (PST) during various stages of harmful algal blooms (HAB) [5–7]. Within the phycosphere interface, exopolysaccharides (EPS) produced by bacterial consortia are the essential intermedia during their dynamic interactions [1,8–10].

*Mameliella* is well-known as one key members belonging to *Roseobacter* group in the family *Rhodobacteraceae* which accounts for a large proportions in bacterioplankton communities in the oceans [11,12]. *Mameliella* spp. are widely contributed to the marine environments in degradation of aromatic compounds, taking part in the biogeochemical cycles of carbon and sulfur, dimethylsulfoniopropionate (DMSP) demethylation, and production of diverse secondary metabolites [13]. The genus *Mameliella* was initially established in 2010 with *Mameliella alba* JLT354-W^T^ as the type species [14]. Previously, four strains, *M. phaeodactyli* and *M. atlantica*, *Ponticoccus lacteus* and *Alkalimicrobium pacificum* were reclassified as later heterotypic synonyms of *M. alba* based on phylogenomics analysis [15].

To convey the composition of microbial consortium governing the phycosphere microbiota (PM) that are closely-associated with diverse HAB dinoflagellates, we have performed the Phycosphere Microbiome Project (PMP) [7–10,17–19]. During the investigation, we found that *Mameliella* is one predominant group and widely distributed in PM of various HAB dinoflagellates analyzed by metagenomics study (Fig. S1). Consequently, during our microbial cultivation study on diverse PM, a new versatile, rod-shaped, halophilic and red-pigmented bacterium designated as strain LZ-28 was isolated from highly-toxic HAB *Alexandrium catenella* LZT09. The new isolate produces active bioflocculanting EPS screened by our flocculanting assay [8–10]. Besides, it shows obvious microalgae growth-promoting ability to its host. Thus, the present work is to clarify the taxonomic classification of this new isolate by both taxonogenomics and phylogenomics approaches along with phenotypic and genotypic characterizations. Furthermore, this study is trying to throw light on the mechanisms governing the algae-bacteria interactions of this novel member of *Mameliella alba* by deeply digging the comparative genomics of *Mameliella* strains.

## Materials and methods

### Bacterial strain and growth media

Strain LZ-28 was isolated from toxic *A. catenella* LZT09 according to our protocol previously described [8–10]. The strain was purified and maintained on marine agar (MA; Difco), and preserved as a glycerol suspension (20%, v/v) at −80 °C for long term preservation, and was transferred into marine broth (MB) for experimental needs.

### Morpho-physiological and biochemical characterization

#### Microscopy observation

Strain LZ-28, grow in exponential phase, was observed its vegetative cells and spores under transmission electron microscopy (JEM-1200; JEOL, Tokyo, Japan) and light microscope (BH-2; Olympus).

### Physiological and biochemical tests

Gram-staining test was followed by the our procedures described previously [8–10]. Hanging-drop technique was used for observing motility during the exponential phase of culture growth. The growth conditions of temperature was tested using MA medium at the temperature range of 4, 10, 15, 20, 25, 28, 30, 33, 37, 40, 45, 50 and 55 °C. The optimal pH was investigated at different pH, ranging from 4.0 to 11.0, at 0.5 unit intervals, culturing on MB medium at 25 °C for 2 days. Salt tolerance was determined by growing in MB medium containing various NaCl concentrations (0–15.0%, w/v, with interval of 0.5%) while removed the original NaCl. Catalase activity and oxidase activity were detected by adding 3% (v/v) H_2_O_2_ and by using oxidase reagent (bioMerieux), respectively. Anaerobic growth was assessed after 1 week cultivation in an anaerobic chamber (Bactron EZ-2; Shellab) on MA at 28 °C. Tests of phenotypic and enzymic characterizations were carried out using API 20NE, API 20E and API ZYM strips (bioMerieux) according to the manufacturer’s instructions.

### Analysis of bacterial fatty acid profiles

Extraction and identification of cellular fatty acids were performed by standard MIDI System (Sherlock version 6.1; database TSBA6) [8–10]. The respiratory quinones were extracted using chloroform/methanol 167 (2:1, v/v) and identified using an Agilent 1100 HPLC system. The polar lipids were examined by two-dimensional TLC and identified with the method described by Minnikin *et al*. [20]. Analysis of the polar lipids was performed by two-dimensional TLC and then followed by the method of Yang *et al*. [18–20].

### HPLC and LC-MS analysis of carotenoids

The carotenoids produced by strain LZ-28 were extracted and determined as described previously [21] by an Agilent 1200 high performance liquid chromatography (Agilent, Palo Alto, CA, USA) equipped with a diode-array detector (DAD). Separation was carried out on an Agilent RP ODS C_18_ analytical column (5 μm, 4.6 mm x 250 mm I.D.) maintained at 24°C. The Agilent 3D ChemStation software (Version 10.0.07) was used for the instrument control and data analysis. The mobile phase consisted of acetonitrile: dichloromethane: methanol (85:10:5, v/v/v). The flow rate was 1.0 mL min^−1^ and UV-visible DAD detection was monitored at 450 nm. A 20 μL loop was used for injection.

The LC-MS analysis of carotenoids was performed using a Finnigan LCQ^TM^ Deca XP MAX ion trap mass spectrometer (Thermo Finnigan, CA, USA) equipped with an atmospheric pressure chemical ionization (APCI) interface (Finnigan), which connected with a Finnigan Surveyor HPLC system equipped with a quaternary pump, a vacuum membrane degasser, a LightPipe™ UV-visible photodiode-array (PDA) detector, an autosampler. Chromatography was carried out on a column with a 2.1 mm ×5.0 mm I.D. guard column. A gradient consisting of solvent A: 85% acetonitrile, 10% dichloromethane and 5% methanol; solvent B: water, 0-12 min 100% A; 12-35 min 85% A was used. Selected [M+H]^+^ was analyzed by collision-induced dissociation with argon gas, and product ion spectra were recorded.

### Bacterial EPS bioflocculanting and MGP bioactivity assays

Extraction of bacterial EPS produced by strain LZ-28 and bioflocculating activity evaluation were performed according to our procedures reported previously (Duan et al. 2020; Mu et al 2019; Zhang et al 2021). The prepared EPS was dissolved in distilled water for furthering bioflocculation activity (BFC) assay [8–10]. Briefly, the measures using kaolin clay suspension flocculation (KCSF) assay calculated as flocculation rate (FR) were used and performed at a 96-well microplate mode with at least triplicates. Microalgae growth-promoting (MGP) activity of strain AM1-D1^T^ toward *A. minutum* amtk4 in a co-culture system was performed as reported previously [22].

### Phylogenetic analysis

The related 16S rRNA gene sequences of species in the genus *Mameliella* and type strains within the *Rhodobacteraceae* family were retrieved from NCBI, except three strains of PBVC088 and Ep20 of *Mameliella alba*, which were extracted from their genome sequences by PhyloSuite version 1.2.1 [23]. Phylogenetic evolutionary analyse based on 16S rRNA gene sequence were performed using the software package MEGA 7.0. The evolutionary distance was calculated with 1,000 bootstrap replicates.

### Phylogenomic analysis

The draft genome sequence of LZ-28 was obtained from our previous GPM Project [7]. The reference genome sequence (10 *Mameliella* strains, six strains of *Rhodobacteraceae* family and one strain of *Rhodococcus* genus) were retrieved from NCBI. The average nucleotide identity (ANI) analyses and the average amino acid Identity (AAI) were performed using the ANI/AAIMatrix Genome-based distance matrix calculator (http://enve-omics.ce.gatech.edu/g-matrix) and the digital DNA-DNA hybridization (dDDH) analyses were using Genome-to-Genome Distance Calculator (GGDC 2.1) (http://ggdc.dsmz.de/) using Formula 2. The phylogenetic tree based on the 92 bacterial core genes was constructed by using the UBCG pipeline [24]. Further visualization and editing of the UBCG tree was accomplished by EvolView tool (https://www.evolgenius.info/evolview). Phylogenetic trees based on pan genes and 4,472 bacterial core genes were conducted by the bacterial pan genome analysis (BPGA) pipeline v.1.3 using USEARCH with default settings [25].

### Pan-genome characterization

BPGA is used to automate the complete pan-genome study and downstream analyses of prokaryotic sequences [25]. All 11 protein sequences annotated by Prokka [26] were used for the BPGA (sequence identity ≥50%; and E value ≤1e−5) tool to perform core-pan-genome analysis. The gene accumulation curve was generated by the BPGA’s result via gnuplot. Core, accessory and unique genes were classified from the orthologous groups through the USEARCH clustering algorithm [25].

### Comparative function genes analysis

The online COG and KEGG databases accessing via BPGA were used to analyze the function of genes. Core, accessory and unique genes were sorted out, and major concerns were on core and unique genes. The gene clusters were identified by anti-SMASH 4.0 from the reference genome sequences of *M. alba* KD53 [27]. All 11 protein sequences produced by Prokka were used for the eggNOG-mapper to get an overview of the functional genes [28]. All results were manually checked.

## Results

### Phenotypic and biochemical characteristics of strain LZ-28

Strain LZ-28 was isolated from algal cells with an exponentially growing culture of toxic *A. catenella* LZT09. Similar to the other bacterial strains belong to *Mameliella alba*, colonies of train LZ-28 grew on MA are circular, smooth, convex, and light red in color. Cells of strain LZ-28 were Gram-stain-negative, rod-shaped (0.7-1.0 μm width and 2.0-2.9 μm length), and non-motile. Poly-β-hydroxybutyrate (PHB) granules were accumulated at the edge of the cells under transmission electron microscopy observation (Fig. 1). The physiological examinations revealed that strain LZ-28 grew at pH ranging from 5.0 to 10.0, with optimum growth at pH 7.0. Growth temperature ranged from 15 °C to 40 °C with the optimum value of 25 °C at the presence of 1-10% (w/v) NaCl with the optimum of 2.5% (w/v). No growth was detected under anaerobic conditions on MA even after three week-long incubation. Hydrolysis of starch, L-tyrosine, and gelatin was observed under experimental condition. Nitrate was reduced to nitrite, but the reduction of nitrite to nitrogen was not observed. Chemotaxonomic analysis showed that the cellular fatty acids profile of strain LZ-28 consisted of C_18 : 0_, C_18 : 1_*ω*7c 11-methyl and C_19 : 0_ cyclo *ω*8c as major fatty acid profiles. Detailed comparison of the morphological, biochemical and physiological properties of strain LZ-28 and five representative strains of *Mameliella alba* were summarized in Table 1.

**Fig 1.**
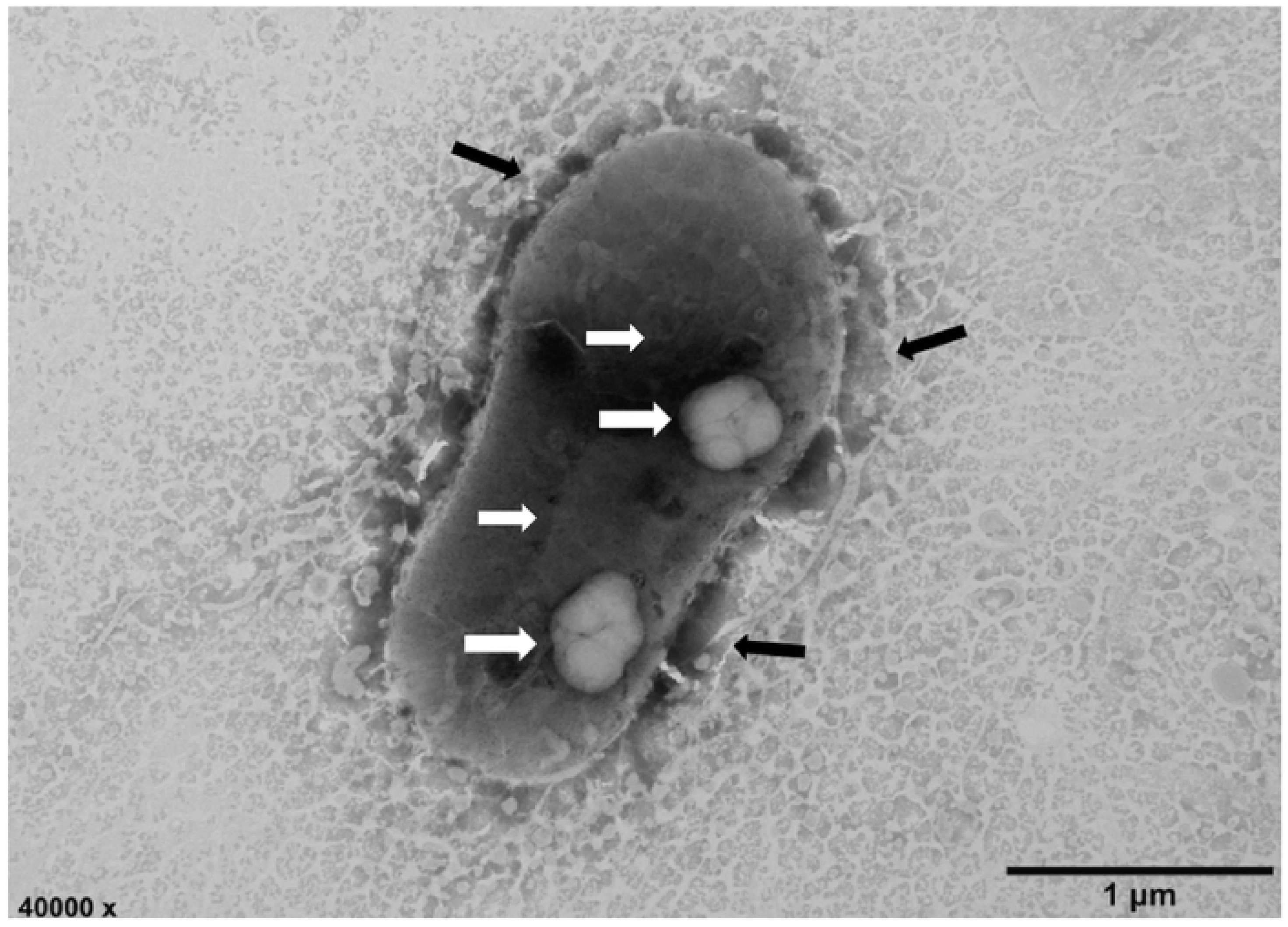
Transmission electron micrograph of the cells of strain LZ-28 grown on MA for 2 days at 25 °C; *Bar* 1 μm. Transmission electron microscopy with white arrow indicating PHA granules, and with black arrow indicating EPS outside the cells

**Table 1.** The morphological and biochemical characteristics of strains LZ-28, JLT354-WT, KD53, L6M1-5, JL351 and F15

| Characteristic | LZ-28 | JLT354-W<sup>Ta</sup> | KD53<sup>a</sup> | L6M1-5<sup>a</sup> | JL351<sup>a</sup> | F15<sup>a</sup> |
| --- | --- | --- | --- | --- | --- | --- |
| Isolation source | Algal<br>phycosphere | Seawater | Algal<br>phycosphere | Deep-sea<br>sediment | Surface<br>seawater | Deep-sea<br>sediment |
| Cell size |  |  |  |  |  |  |
| Width (µm) | 0.7-1.0 | 0.7–0.9 | 0.5–1.0 | 0.7–0.8 | 0.5–0.8 | 0.7–1.2 |
| Length (µm) | 2.0-2.9 | 2.0–3.2 | 2.2–2.9 | 1.0–2.0 | 0.7–1.5 | 1.6–3.4 |
| Growth conditions |  |  |  |  |  |  |
| NaCl range (w/v, %) | 1.0-10.0 | 1.0–10.0 | 1.0–6.0 | 0.5–15.0 | 0.5–9.0 | 0–10.0 |
| Optimum NaCl | 2.5 | 1.0–3.0 | 3.0 | 3.0–5.0 | 1.0–4.0 | 3.5 |
| pH range | 5.0-10.0 | 6.0–9.0 | 6.0–9.5 | 5.0–10.5 | 7.0–10.0 | 6.0–11.0 |
| Optimum pH | 7.0 | 8.0 | 7.5–8.0 | 7.0 | 8.0 | 7.0–8.0 |
| Temperature range (°C) | 15-40 | 10–30 | 16–37 | 10–41 | 15–40 | 4–50 |
| Optimum temperature | 25 | 25 | 28 | 28–30 | 30 | 35–37 |
| Oxidase | + | + | + | – | + | + |
| Catalase | + | + | – | w | + | + |
| Poly-β-hydroxybutyrate | + | + | – | – | + | – |
| API 20E test |  |  |  |  |  |  |
| Citrate utilization | – | – | – | w | w | + |
| Gelatinase | + | + | – | + | + | + |
| API 20NE test |  |  |  |  |  |  |
| Reduction of nitrite | + | + | + | + | – | + |
| Gelatin hydrolysis | + | + | – | + | + | + |
| Utilization of phenylacetic acid | – | + | + | + | + | – |
| Polar lipids |  |  |  |  |  |  |
| Phosphatidylcholine (PC) | + | – | + | – | + | + |
| Diphosphatidylglycerol (DPG) | + | – | – | – | + | – |
| Phosphatidylethanolamine<br>(PE) | + | – | + | + | + | + |
| Aminolipids (AL) | – | + | + | + | + | – |
| Glycolipid (GL) | – | – | – | – | + | + |
| Fatty acids profiles |  |  |  |  |  |  |
| C<sub>16:0</sub> | 3.2 | 5.7 | 5.3 | 4.8 | 5.3 | 4.5 |
| C<sub>17:0</sub> | 1.1 | TR | 1.0 | TR | TR | ND |
| C<sub>18:0</sub> | 7.0 | 9.2 | 11.1 | 10.0 | 8.6 | 8.5 |
| C<sub>12:1</sub> 3OH | 4.7 | 3.3 | 2.8 | 3.1 | 3.2 | 3.1 |
| C<sub>17:1</sub>ω8c | – | 0.9 | TR | 0.7 | 0.6 | 1.0 |
| C<sub>18:1</sub>ω7c 11-methyl | 8.7 | 3.7 | 5.6 | 4.7 | 4.1 | 4.3 |
| C<sub>19:0</sub> cyclo ω8c | 7.7 | 3.7 | 1.7 | 2.9 | 3.2 | 3.2 |
| Summed feature 8* | 65.5 | 70.8 | 69.5 | 70.7 | 72.5 | 74.3 |
| 16S rRNA similarity to LZ-28 | 100.00 | 99.70 | 99.77 | 99.92 | 99.62 | 99.40 |
| DNA G+C content (mol%) | 64.9 | 65.2 | 65.0 | 65.0 | 65.1 | 65.0 |
| ANI value to LZ-28 (%) | 100.0 | 98.0 | 100.0 | 98.0 | 98.0 | 97.9 |
| AAI value to LZ-28 (%) | 100.0 | 98.4 | 100.0 | 98.1 | 98.5 | 98.2 |
| dDDH value to LZ-28 (%) | 100.0 | 84.3 | 100.0 | 83.9 | 84.3 | 83.1 |

+, positive; W, weakly positive; –, negative.
\* Summed feature 8 comprises C<sub>18:1</sub>ω7c and/or C<sub>18:1</sub>ω6c.
TR, trace amount (less than 0.5%). ND, not detected.
Data from the original study: <sup>a</sup>, Liu *et al.* (2018).

### 16S rRNA phylogenetic analysis of *M. alba* strains

In the constructed phylogenetic neighbor-joining (NJ) tree based on 16S rRNA gene sequences, strain LZ-28 formed a monophyletic clade with type and non-type strains of *M. alba*, compared with other members of the family *Rhodobacteraceae* (Fig. 2). Previously, *M. phaeodactyli, M. atlantica, Ponticoccus lacteus* and *Alkalimicrobium pacificum* were reclassified as later heterotypic synonyms of *M. alba* [13]. In this study, the phylogenetic analysis based on 16S rRNA gene sequences showed that strain LZ-28 shares high 16S rRNA gene similarity values of 99.70%, 99.77%, 99.92%, 99.60% and 99.40%, with strains of JLT354-W^T^, KD53, L6M1-5, JL351 and F15, respectively, although they were obviously isolated from different sources (Table S1). These values are all exceeding the threshold values (98.65%) of 16S rRNA gene generally accepted for species delineation (Table 1). This phylogenetic characterization was also confirmed by two other phylogenetic trees constructed by maximum-likelihood (ML) (Fig. S2) and maximum-parsimony (MP) algorithms (Fig. S3), respectively, via bootstrap analysis with 1,000 replications. These result clearly indicated that strain LZ-28 is affiliated to the genus of *Mameliella*, and probable a new member of *M. alba*.

**Fig 2.**
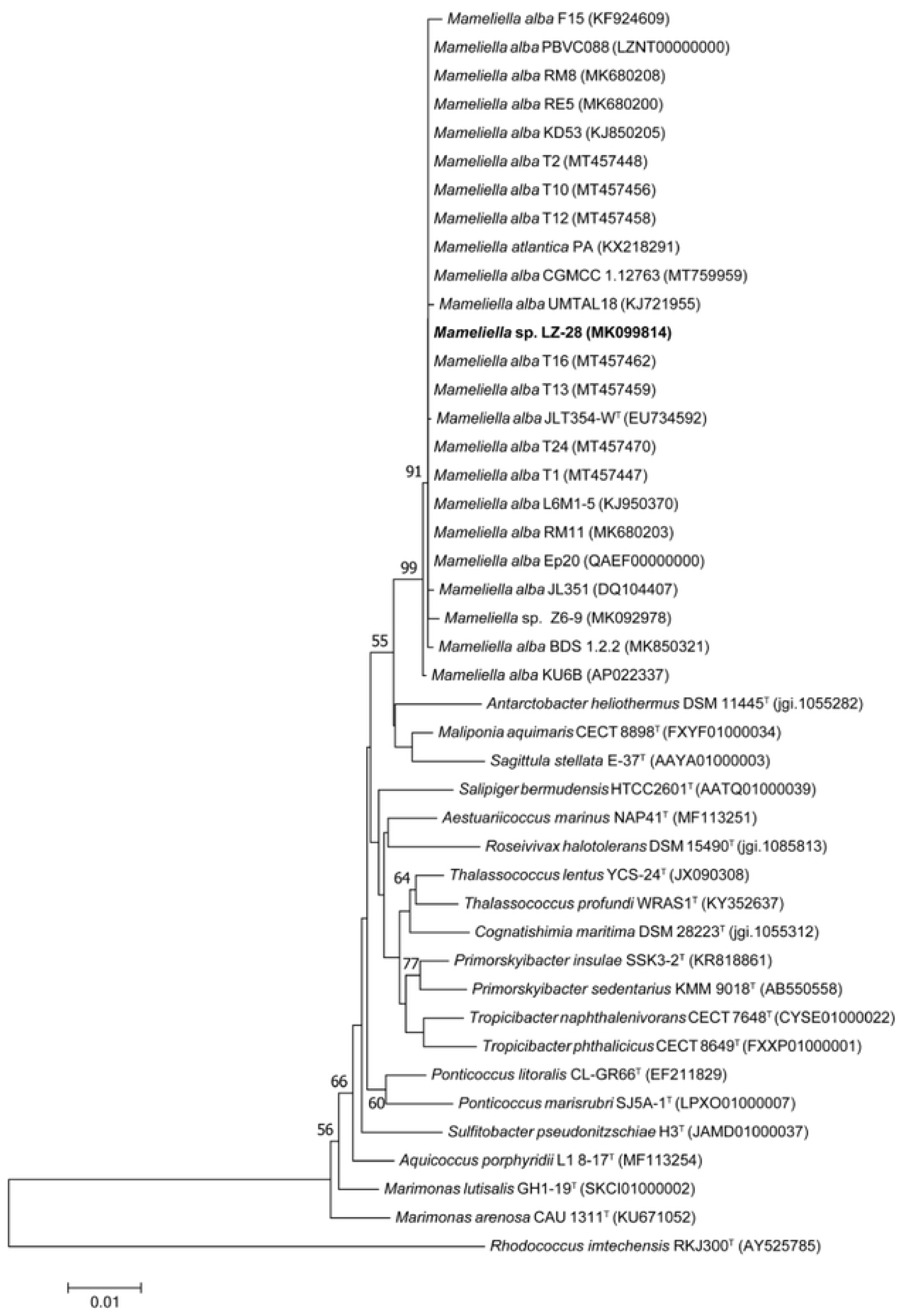
Phylogenetic tree (NJ) based on 16S rRNA gene sequences showing the relationship between strains of the genus *Mameliella* and type strains within the family *Rhodobacteraceae*. Bootstrap values (≥ 50%) based on 1000 replications are shown at the branch points. *Bar*, 0.01 substitutions per site. *Rhodococcus imtechensis* RKJ300^T^ was used as an outgroup

### Genomic features of *M. alba* strains

Due to the high 16S rRNA gene sequence similarities of the new isolate, the phylogenomic analysis was then performed. The genomes of 9 *M. alba* strains were obtained from NCBI (https://www.ncbi.nlm.nih.gov) along with the genome of strain LZ-28. Among the 10 *M. alba* strains, the 9 strains including strains LZ-28 were draft genomically sequenced and strain KU6B has a complete genome. The general genomic characteristics of the 10 strains are summarized in Table S1. The genomic size of strain LZ-28 was 5.66 Mb with G+C content of 64.9%. It contains 5,376 proteins and 58 RNA genes including 8 rRNA and 50 tRNA. And the *N*_50_ for the 39 contigs were 326,544 bp. Genome of strain LZ-28 is the 3^rd^ largest falling into the range of 5.26-5.90 Mb among the 10 *M*. *alba* strains. However, genomic DNA G+C content of strain LZ-28 is the lowest (64.9 mol%) among members of *M*. *alba* strains. Table S1 summarizes several key genomic features for the genomes of 10 *M*. *alba* strains.

### Phylogenomic characterization of strain LZ-28

To further infer the phylogenetic relationship among all *M*. *alba* strains with available genome sequences, the genome-based phylogenetic tree using an up-to-date bacterial core gene set (UBCG) consisting 92 bacterial core genes was performed. The obtained result was shown in Fig. 3. It clearly shows that strain LZ-28 clusters together with strain KD53, which was isolated from marine diatom *Phaeodactylum tricornutum* [13]. Phylogenetic trees constructed by bacterial pan genes (Fig. S4) and 4,472 bacterial core genes (Fig. S5) showed little difference of phylogenomic relationship among 10 *M. alba* strains in the constructed UBCG tree. Additionally, as shown in Fig. 4, the comparison of the ANI (pane A), AAI (pane B) and digital DNA-DNA hybridization (dDDH, in pane C) values of strain LZ-28 and the type strain of *M. alba* JLT354-W^T^ were 98.0%, 98.4%, and 84.3%, respectively, both higher than the 95-96%, 96% and 70% identity thresholds for ANI, AAI and dDDH. Accordingly, it taxonogenomically confirmed that strain LZ-28 is a novel member of *M. alba*.

**Fig 3.**
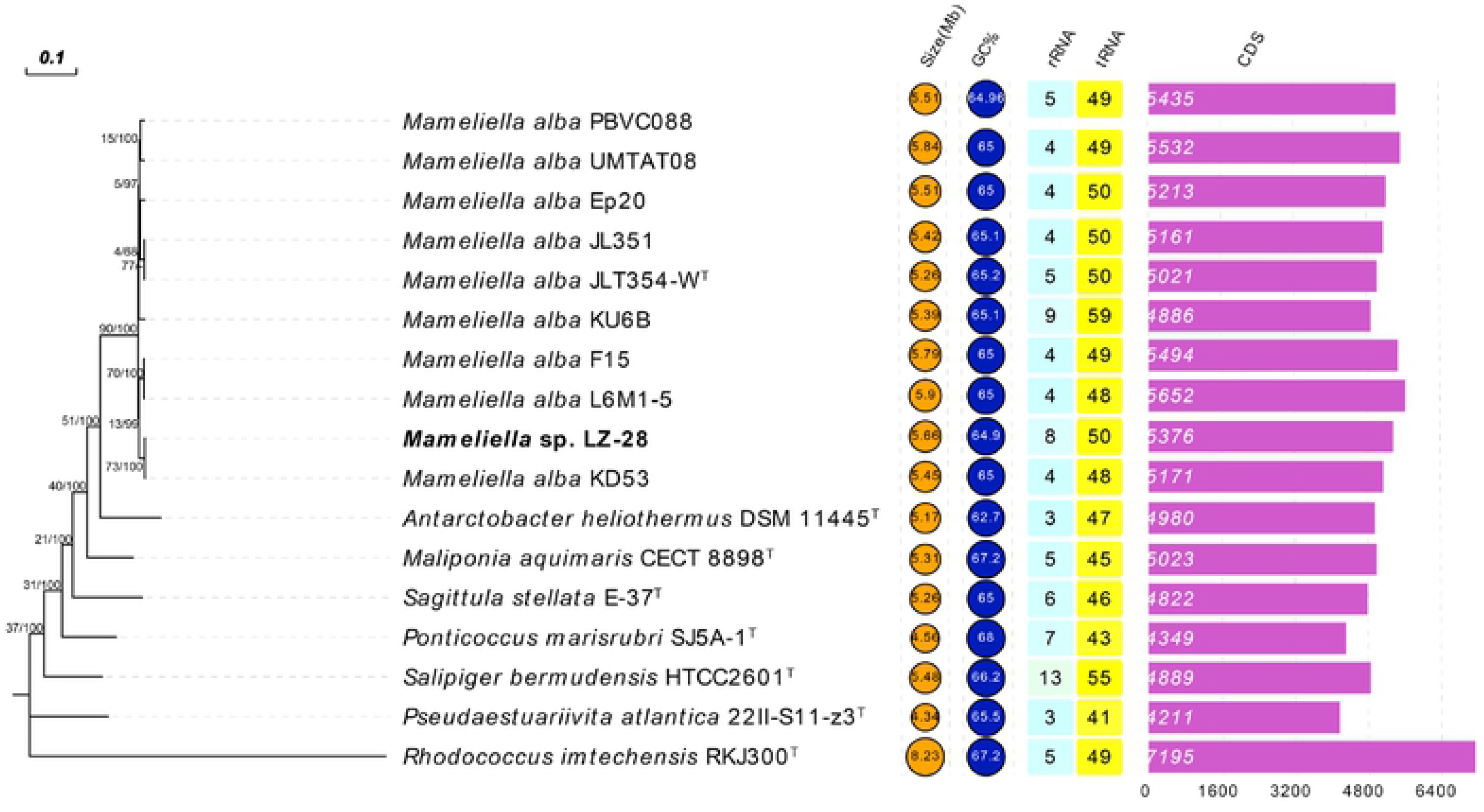
Phylogenomic tree based on 92 bacterial core genes revealing the genome correlation between genomes of the 10 strains of *M. alba* and type species within the family *Rhodobacteraceae*. Gene support index (the number of individual gene trees presented the same node in total genes used) (left) and bootstrap values (right) are given at nodes. GenBank accession numbers are shown in parentheses. *Bar*, 0.10 nt substitution rate (Knuc) units

**Fig 4.**
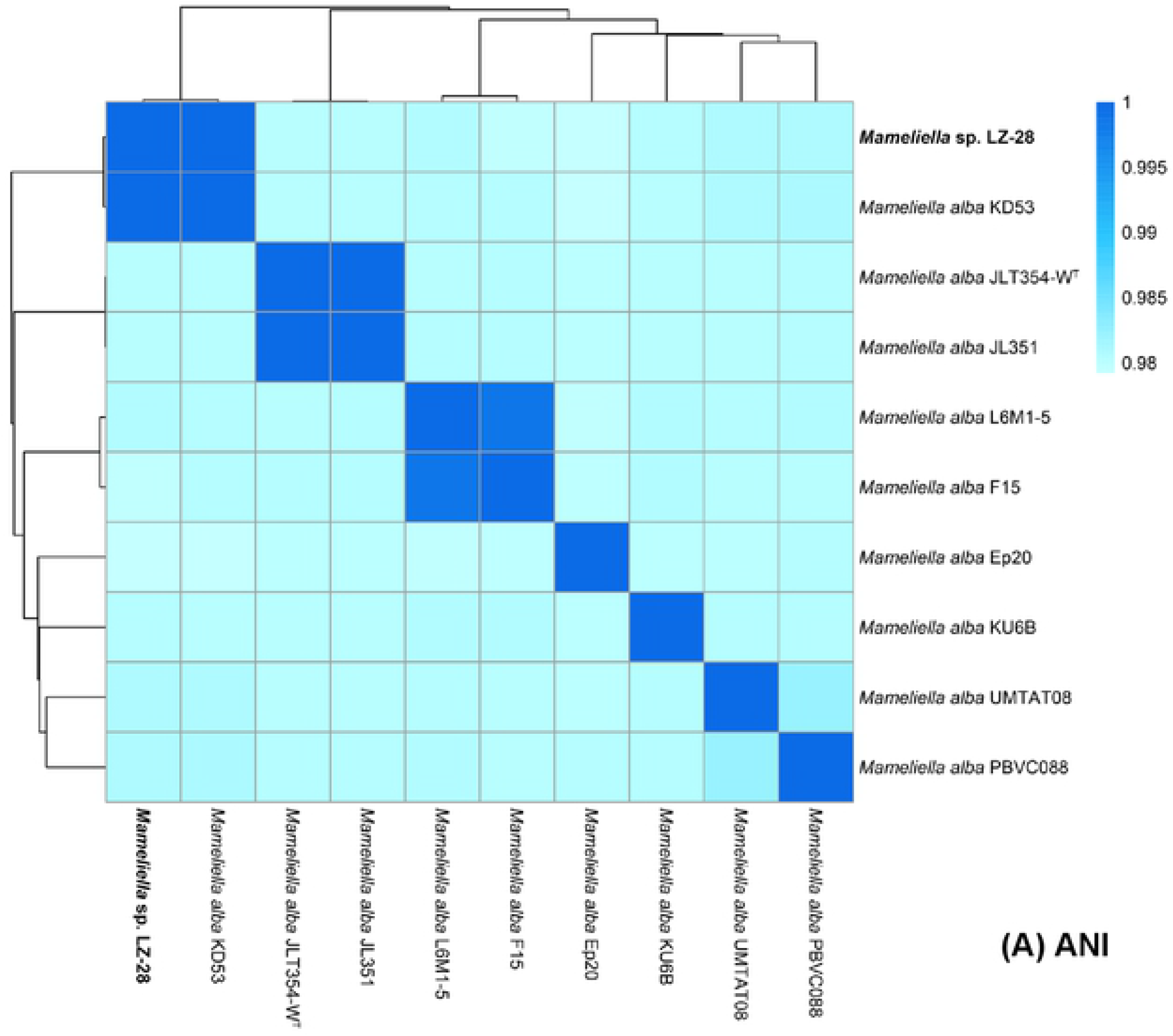
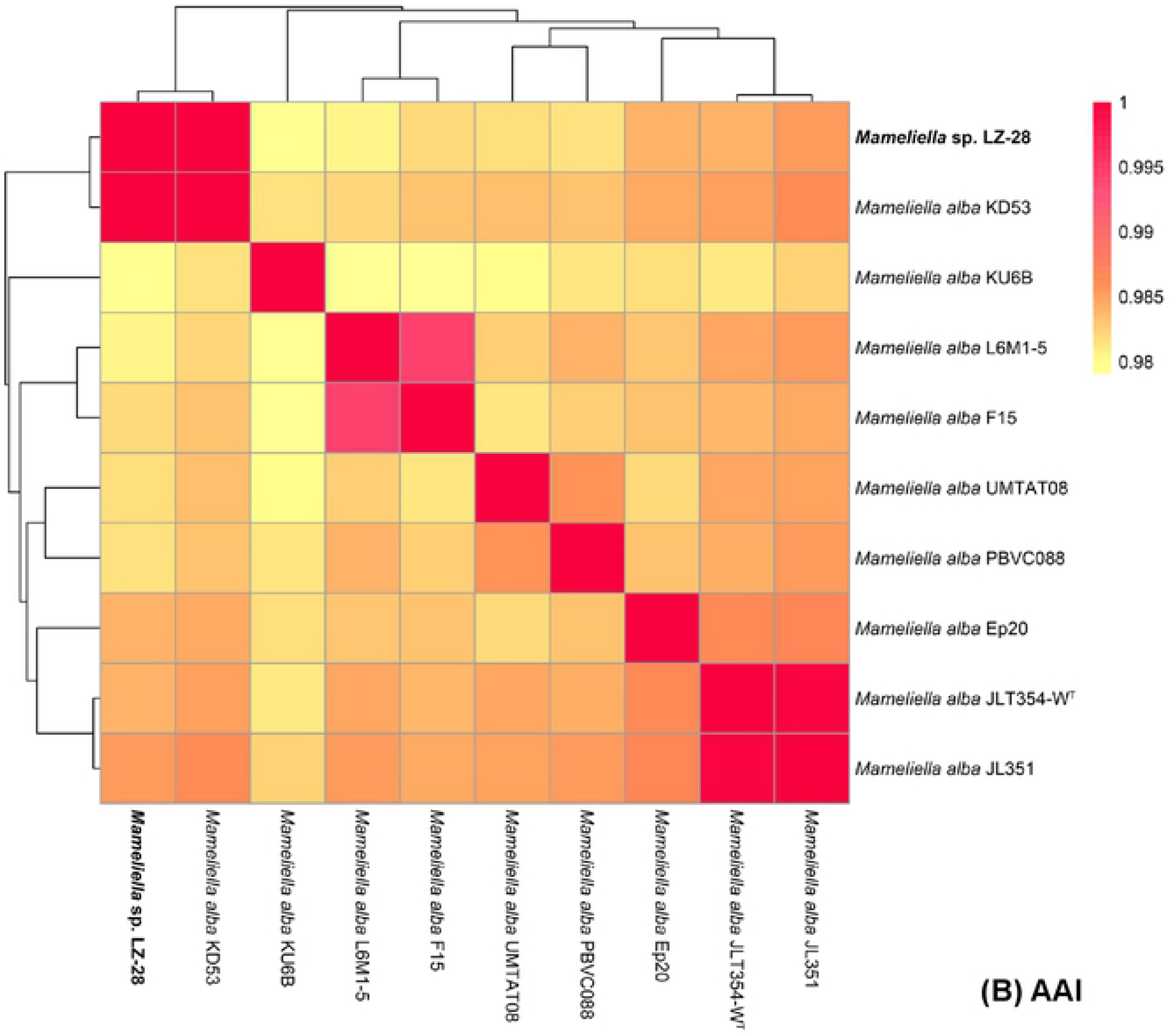
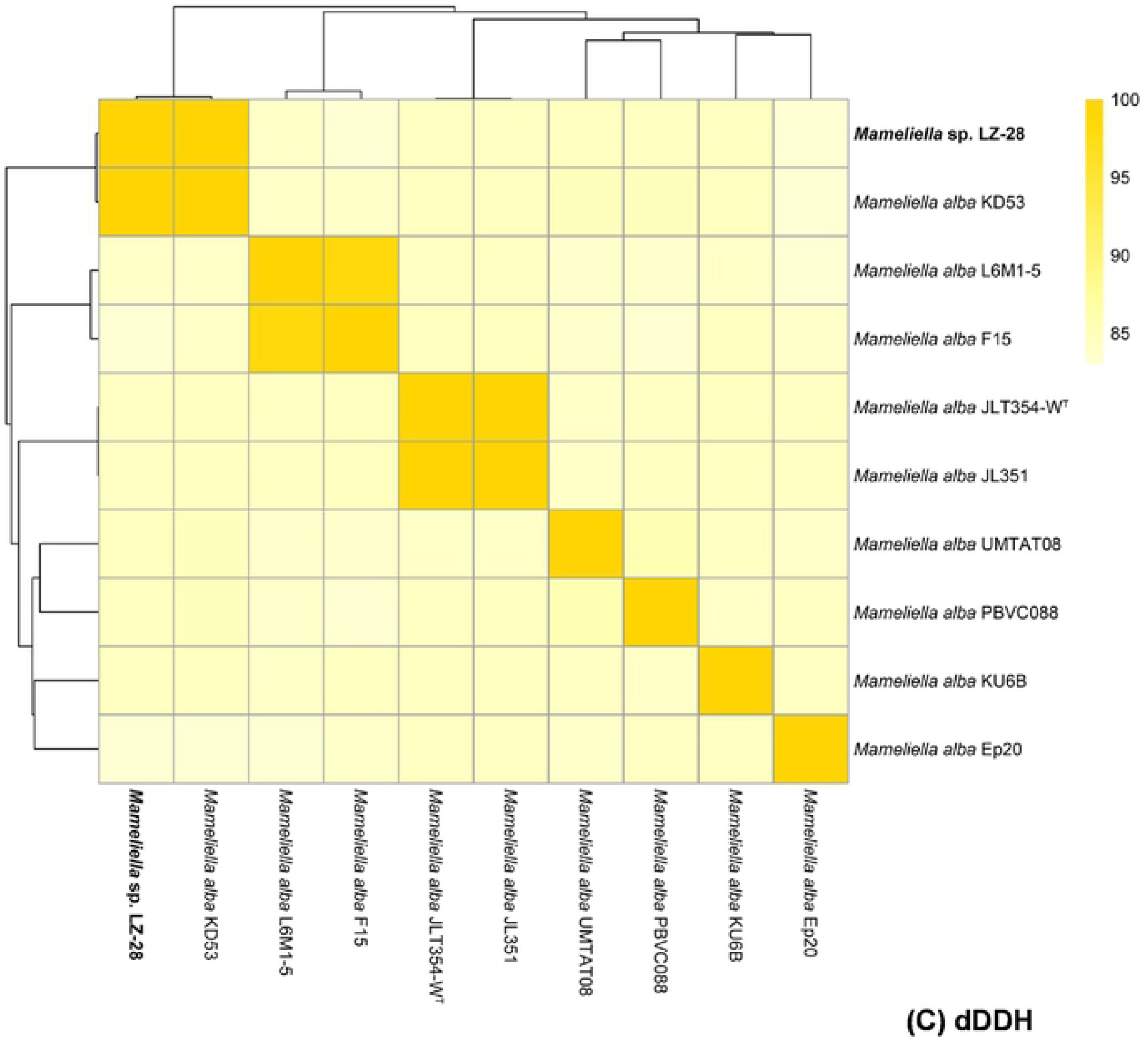
Comparison of the (A) average nucleotide identity (ANI), (B) average amino acid identity (AAI) and (C) dDDH values based on the genomes of 10 strains of *M. alba*.

### Bacterial core- and pan-genomic comparison of *M. alba* strains

The genome sequences of 9 *M. alba* strains were retrieved from NCBI database. Two function curves were constructed according to the core- and pan-genome analyses, respectively. The pan-genome of 10 strains of *M. alba* was fitted into a power law regression function [Fp(x)=5160.29*x^0.15^], indicating that the pan-genome is in an open state. This kind of open pan-genome often exists in bacterial species residing in various ecological environments, and has the tendency of horizontal gene transfer (HGT) [30]. And the core-genome was fitted into an exponential regression [Fc(x)=4965.15*e^−0.01x^] accordingly. From the two curves, the gene number of the pan-genome increased with the strain number of *M. alba* accumulated. Meanwhile, the tendency of core genome was on the contrary, implying that more additional strains would result in more additional unique genes. With the increasing of the gene number as the addition of genome, the pan-genome curve did not reach the plateau. The number of accessory and unique genes of *the* 10 *M. alba* strains were exhibited in Fig. 5. Based on the results, about 4,472 core genes were shared by 10 *M. alba* strains, accounting for 80.8% to 90.3% of the genome repertoire of each strain. And only 0.05% to 5.11% of the unique genes were distributed in individual strain. It indicates that the highly conservative evolution among the members of *M. alba* investigated in this study. The significant difference of the number of unique genes among 10 strains of *M. alba* may be explained by their adaption to growth conditions and possible horizontal gene transfer (HGT).

**Fig 5.**
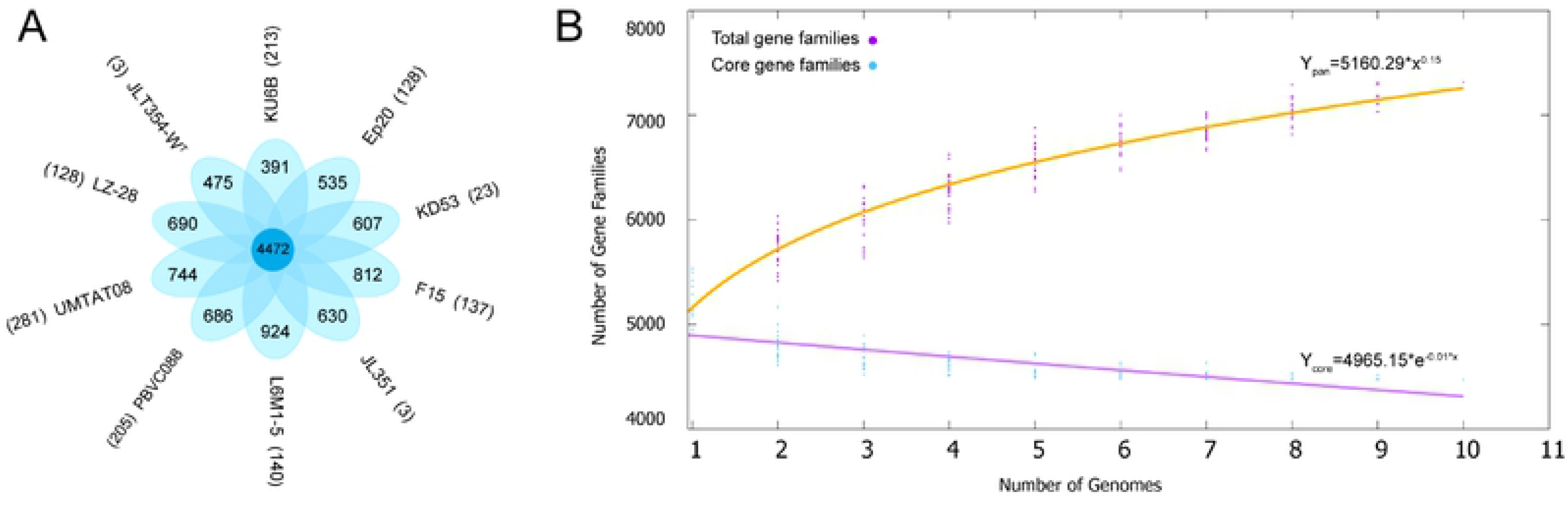
Core–pan-genome comparison of 10 strains of *M. alba* by BPGA analysis Pane A: Petal diagram of the core, accessory, and unique genes of the pangenome. The number of core genes is displayed in the central circle. Overlapping regions represents the number pf accessory genes. Strain names are located outside each petal along with the strain-specific gene number showing in brackets. Pane B: Core-pan-genome accumulation curves of the 11 *M. alba* strains. The purple and blue dot stand for the number of pane genes and core genes respectively.

### Comparison of functional classes of predicted genes of *M. alba* strains

The functions of gene families in genomes of 10 *M. alba* strains was evaluated by performing the COG categories and KEGG categories analyses. The obtained result is presented in Fig. 6. The COG distribution profile showed that most core genes and unique genes were both mainly related to carbohydrate transport and metabolism [G], Transcription [K], Replication, recombination and repair [L], General function prediction only [R] and Function unknown [S]. The essential genes including ABC transporters (*gln*H, *art*Q, *yej*B, *yej*E, *yhd*Y), transcriptional regulators (*cdh*R, *cys*L, *hex*R, *lut*A, *mcb*R, *isc*R and *znt*R) and DNA polymerase (*pol*A, *dna*Q, *dna*X and *dinB*), played a key role in obtaining necessary nutrients from varied environments and maintaining a basic lifestyle for bacteria. In addition, strain UMTAT08 isolated from marine dinoflagellate *Alexandrium tamiyavanichii* harbored the largest core gene numbers during COG analysis. Most unique genes were only found to distribute in strain KU6B isolated from surface seawater in Japan [20]. With respect to the KEGG assignment, genes related to general function, amino acid metabolism and carbohydrate metabolism accounted for the major genes types based on the characterization of the core- and unique-genomes of the 10 *M. alba* strains. However, there was no unique gene sorted in COG categories and KEGG categories in strain JL351 [27]. It may indicate their high conservation of bacterial evolution.

**Fig 6.**
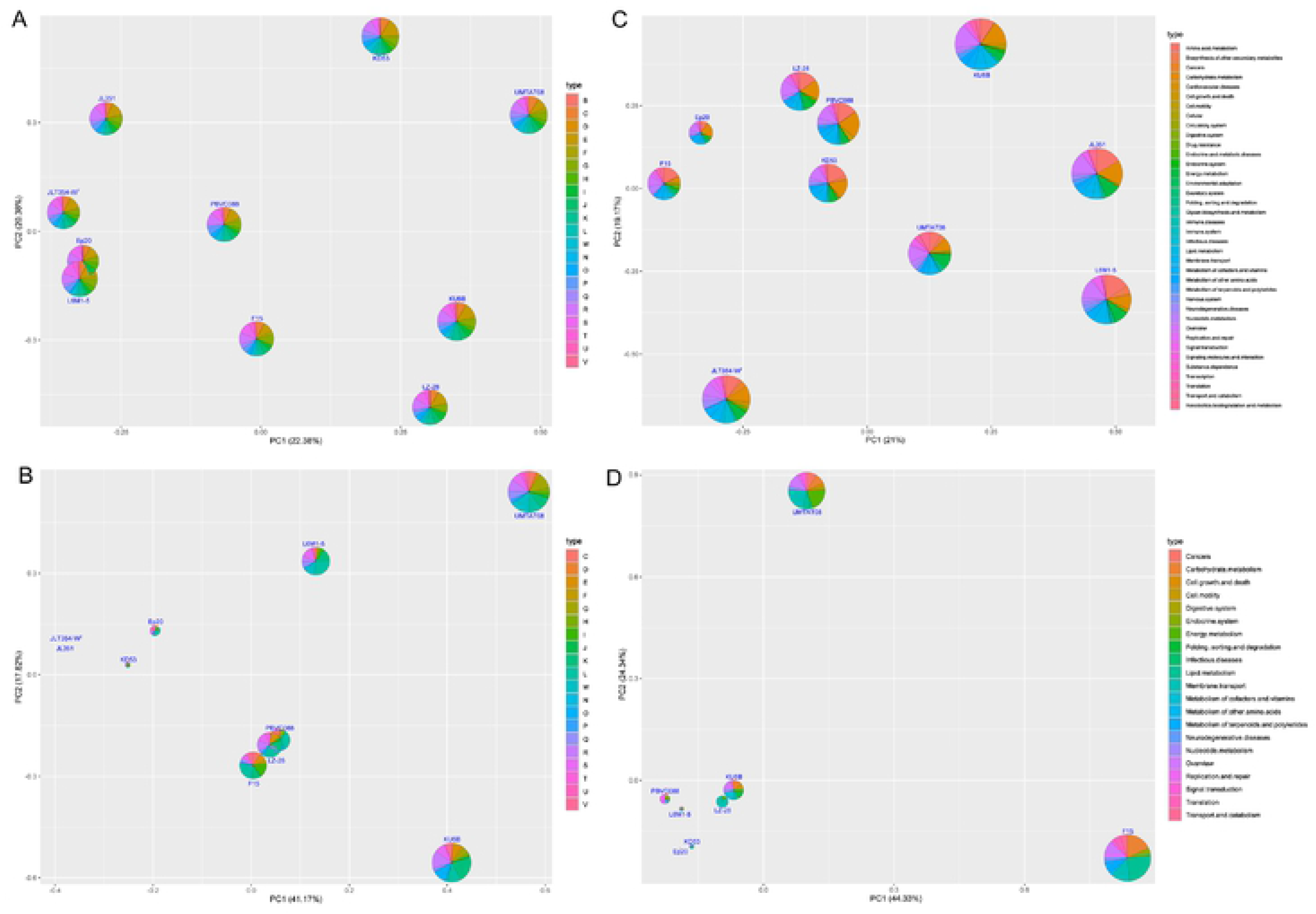
Comparison of COG categories and KEGG categories among genomes of 10 strains of *M. alba* The functional genes of core and unique groups distributed by COG database are shown in the left pane A and pane B, respectively. The panes C and D represent the distribution of core genes and unique genes annotate by KEGG database, respectively.

Genes encoding for polyhydroxyalkanoate (PHA) or PHB, including *pha*A, *pha*B, *pha*C, *pha*D, *pha*E, *pha*J, *pha*P, *pha*R and *pha*Z were found in genomes of some *M. alba* strains. These PHA/PHB biosynthesis genes did not form a gene cluster owing to the long distance between each other. Moreover, genes of *frc*A (fructose import ATP-binding protein *frc*A), *frc*B (fructose import binding protein *frc*B), *frc*C (fructose import permease protein *frc*C), *ytf*T (inner membrane ABC transporter permease protein *ytf*T) and *ytf*Q (ABC transporter periplasmic-binding protein *ytf*Q), which were involved in ABC-type sugar transport system periplasmic component, were all found in the genomes of the 10 *M. alba* strains. Additionally, they all contained aldehyde dehydrogenase family genes (*gab*D2, *gab*D) and aldehyde oxidase and xanthine dehydrogenase, a/b hammerhead domain genes (*xdh*A, *cox*L), performing the function of nutrient and metabolite transport.

In addition, PHA or PHB granules like substances presented in the cytoplasm of strain LZ-28 strain were genomically verified by the presence of PHA biosynthesis genes in the genome. The similar observation was also found in both strains JLT354-W^T^ and JL351 that accumulate PHB granules. Furthermore, predicted genes which were connected with the nutrients and metabolites trafficking (*frc*A, *frc*B, *frc*C, *ytf*T, *ytf*Q, *gab*D2, *gab*D, *xdh*A and *cox*L) and photosynthetic genes (*crt*A, *crt*I, *crt*B, *crt*C, *crt*D, *crt*E and *crt*F) were all found in the 10 strains of *M. alba* genomes

### Characterization of potential algae-bacteria interactions

Several key biosynthetic gene clusters (BGCs) responsible for synthesis of bacterial secondary metabolites were identified in strain LZ-28, and also compared with other *M. alba* strains. The *crt* gene cluster was found in strain LZ-28 including the *crt*A, *crt*B, *crt*C, *crt* D, *crt*E, *crt*F and *crt*I encoding for carotenoid biosynthesis (Fig. 7). To confirm the metabolites of *crt* gene cluster in strain LZ-28, the methanol extracts were detected by HPLC and LC-MS analysis. It exhibited three major peaks (T*_r_* = 7.588, 19.781 and 23.685 min, respectively) and several minor peaks monitored at 450 nm in HPLC profile (Fig. 8). The three main peaks showed typical spectrum of carotenoids, and showed the protonated molecule [M+H]^+^ at *m/z* at 566.6, 589.5 and 540.7 corresponding to the speculated molecular formula of C_40_H_52_O_2_, C_40_H_54_O_3_ and C_40_H_52_O, respectively (Fig. 8), although their detailed chemical structures remain to be elucidated. Carotenoids usually act as key light-gathering pigments and essential ingredients in photosynthesis, also protects bacterial chlorophyll against photooxidation [31]. Moreover, it’s interesting to note that not every *M. alba* strain produce carotenoid pigments although they all have *crt* gene cluster for carotenoid biosynthesis (Table 1). The reason may lie in the expression of the *crt* genes are regulated by the culture conditions or environmental factors [31].

**Fig 7.**
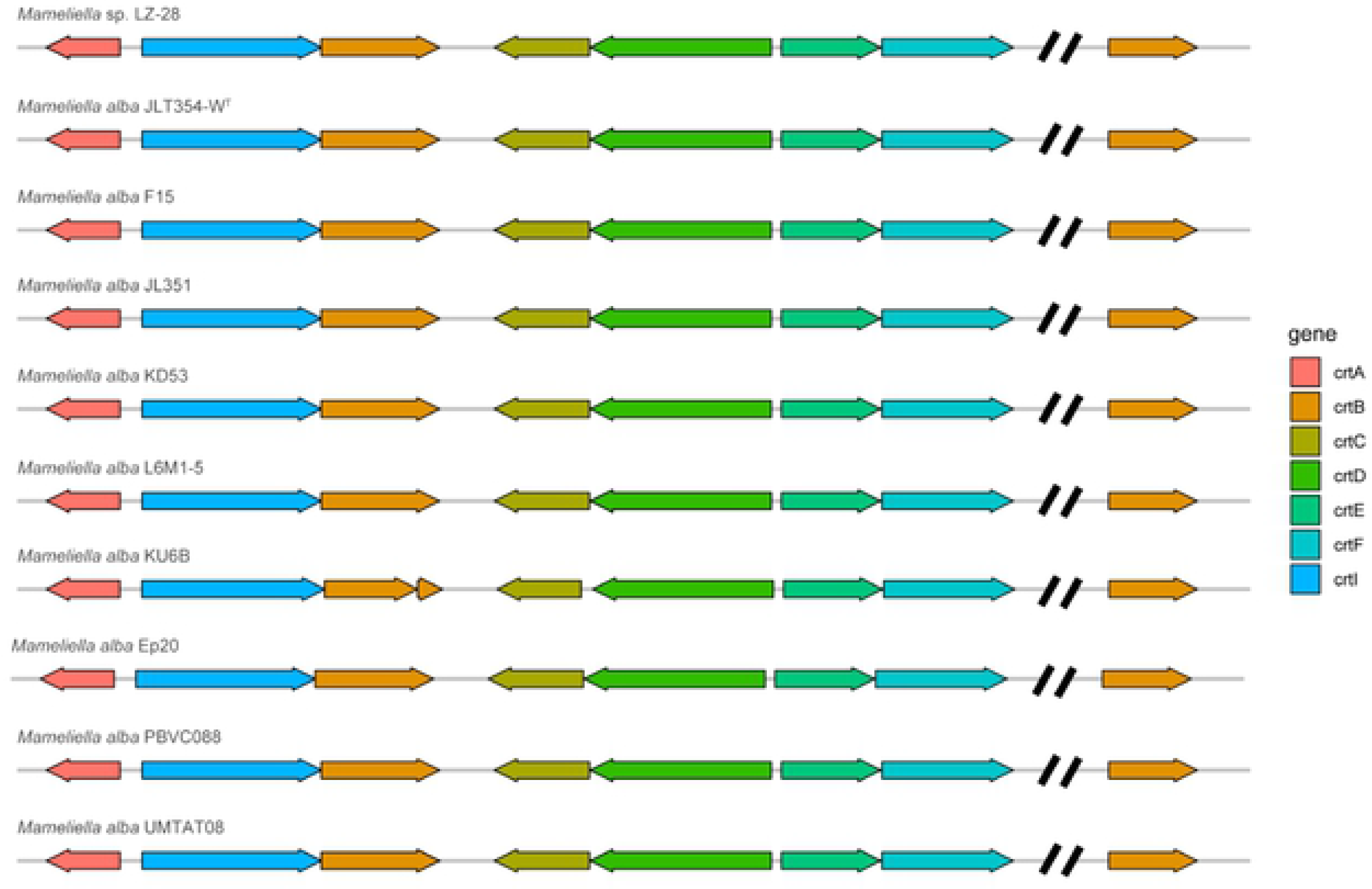
Comparison of biosynthetic gene cluster of carotenoids between the genomes of 10 strains of *M. alba* Different color arrows indicate different genes and functional genes belonging to genome scaffolds are separated by two slashes.

**Fig 8.**
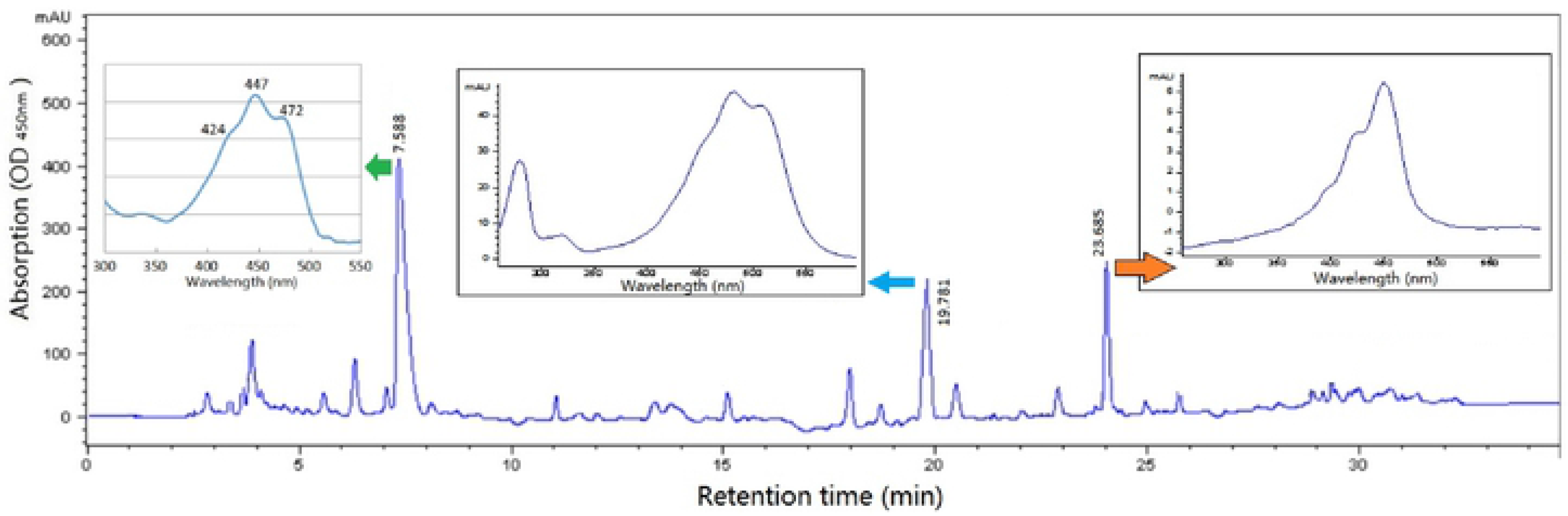
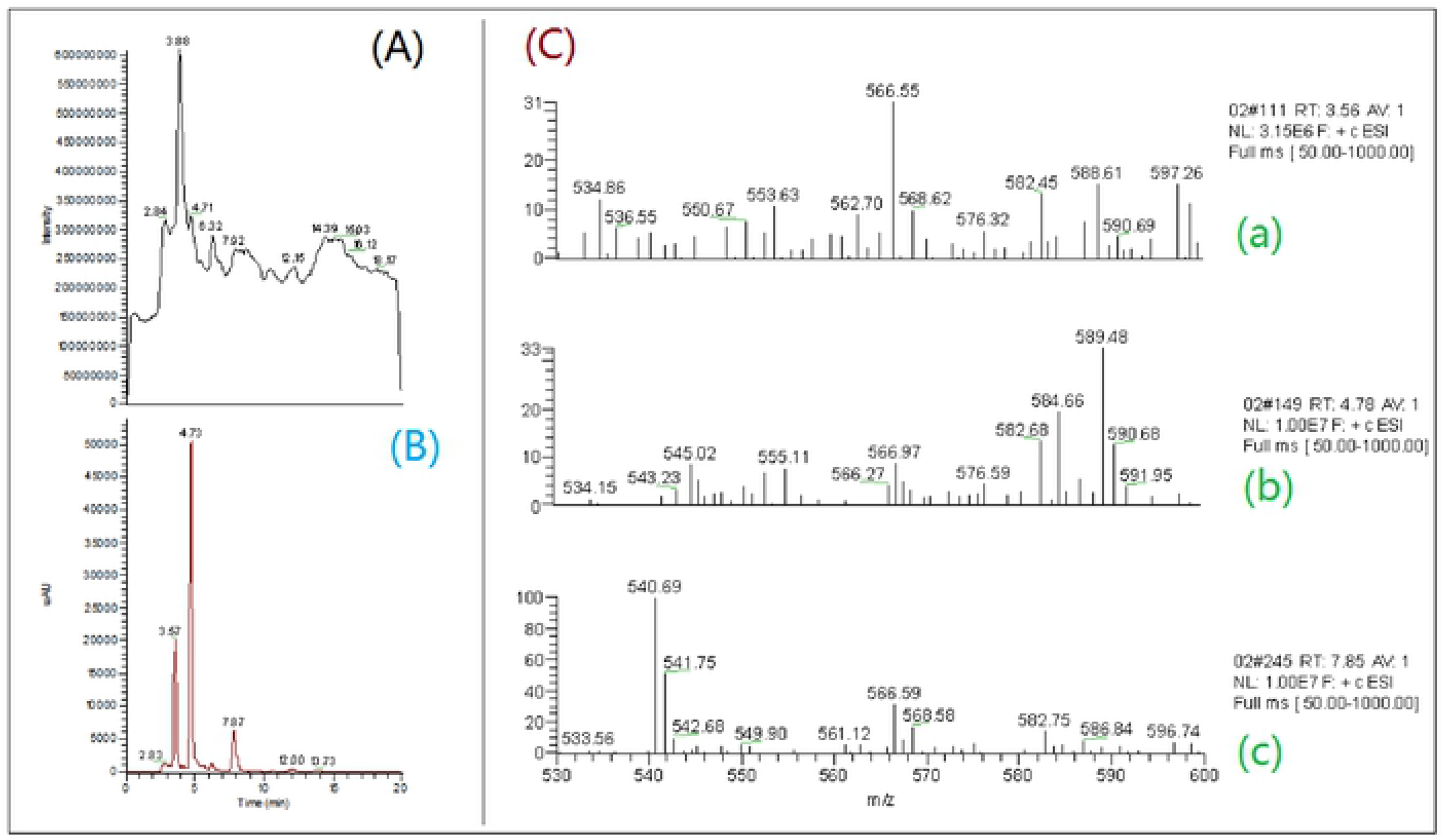
HPLC and LC-MS chromatograms of the carotenoids of strain LZ-28 Conditions were as described in the Materials and methods section.

Most bacteria including strain LZ-28 have enzymes that require vitamin B_12_ (cobalamin) as a cofactor [32]. The genomic mining revealed the complete *cob* gene cluster (*cob* A-Z) encoding for vitamin B_12_ biosynthesis in all 10 *M. alba* strains (Fig. 9). Vitamin B_12_ biosynthesis have been only characterized in prokaryotic organisms. Many algae have been found to require exogenous cobalamin for growth in culture, implying that they are unable to make it themselves. However, algae may acquire vitamin B_12_ through a symbiotic relationship with closely-associated bacteria [33].

**Fig 9.**
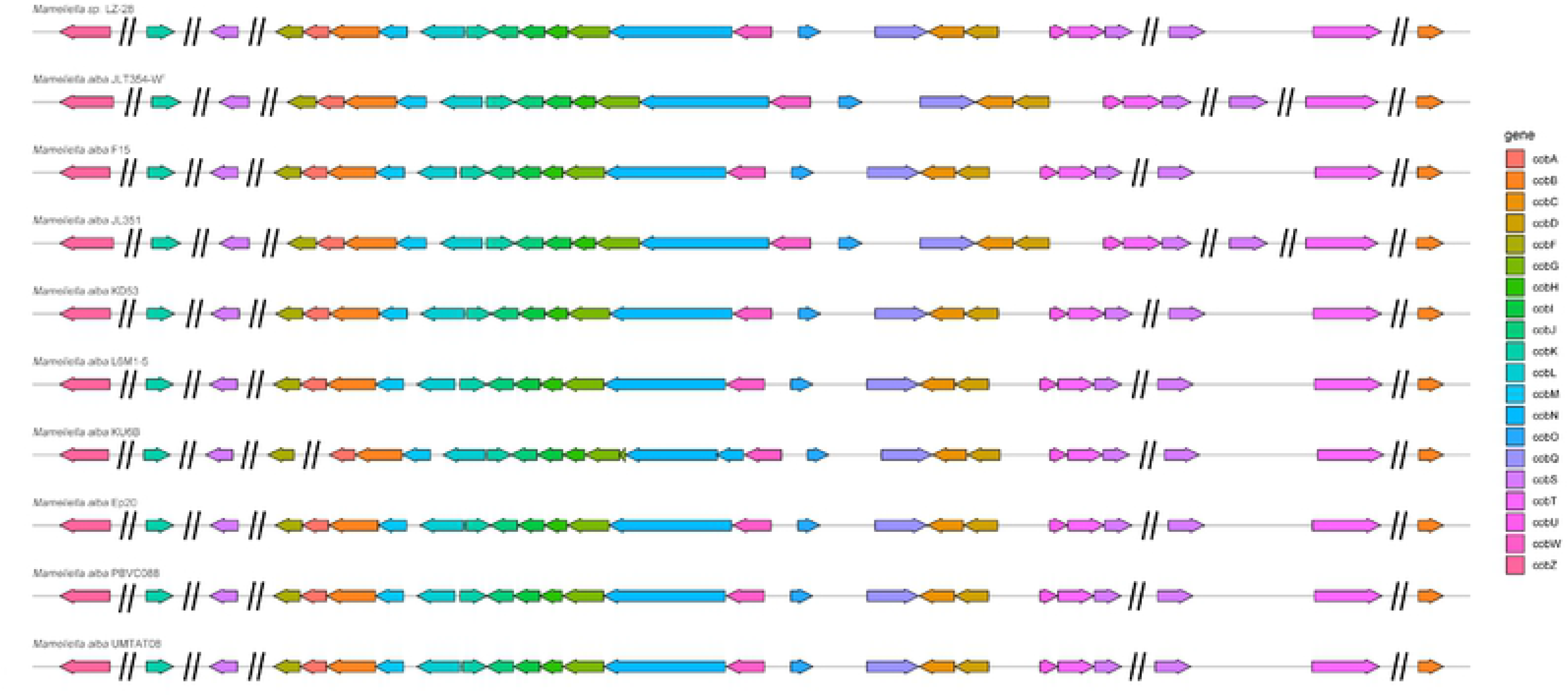
Biosynthetic gene cluster of vitamin B_12_ of the genomes of 10 strains of *M. alba* Different color arrows indicate different genes and the functional genes belonging to different genome scaffolds are separated by two slashes.

The bacterial EPS have been revealed to serve as one vital chemical intermedia within the microscopic phycosphere niches, and they usually mediate the host-microbe interactions [1–3]. Based on the result obtained by our bacterial bioflocculanting (BFC) assay, the EPS of strain LZ-28 demonstrate obvious bioflocculanting activity with concentration-dependent manner (pane A, Fig. 10). It can be seen that the BFC reached the maximum of 95.7±8.5% at 0.60 g L^−1^ of bacterial EPS of strain LZ-28, which exhibiting higher bioflocculanting deficiency compared with the strains reported previously [8–10]. For MGP assay, strain LZ-28 demonstrated obvious microalgae growth-promoting activity when co-cultured with host LZT09 (pane B, Fig. 10). Those results clearly indicate that strain LZ-28 is a new MGP bacterium, and produces novel bioactive bioflocculating EPS with potential environmental and biotechnological implications [8–10]. Also, several biosynthesis genes (*wzx, exo* and *muc*) for bacterial EPS biosynthesis were found in the genome of strain LZ-28. Thus, this study indicated that strain LZ-28 could serve as a novel bacterial candidate with natural potential for the production of promising and versatile microbial bioflocculants derived from the unique marine phycosphere niches.

**Fig 10.**
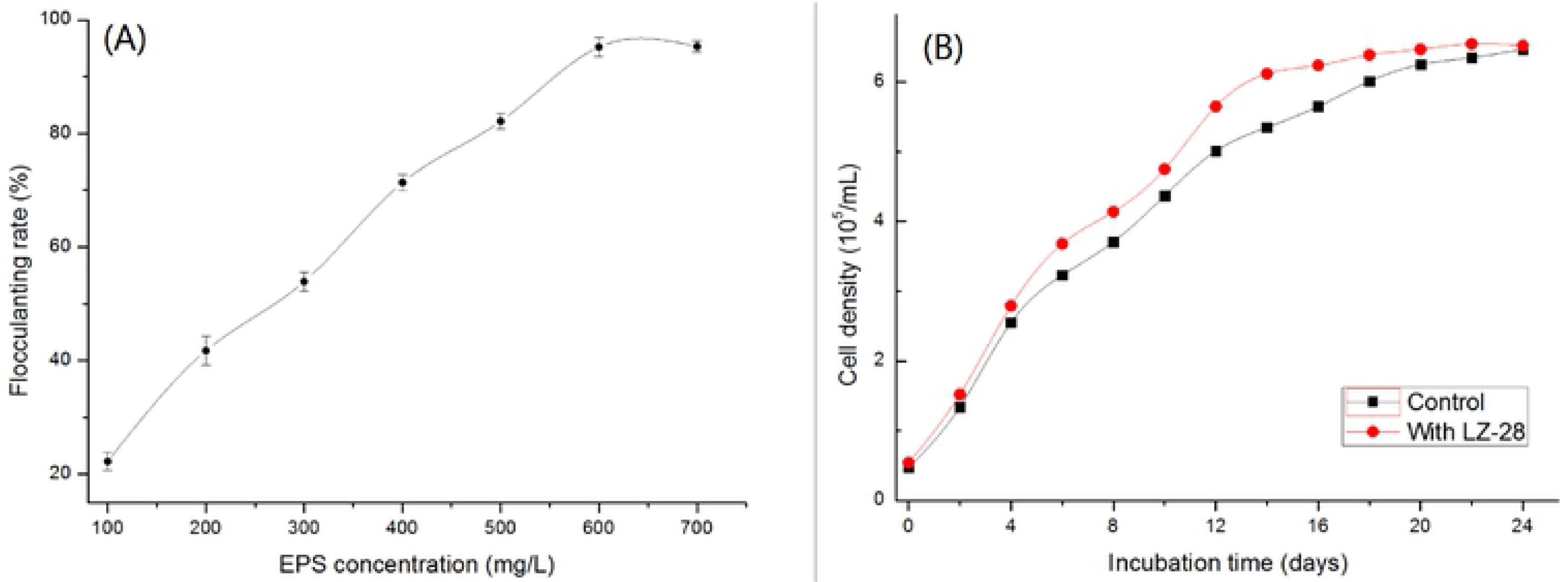
The bioactivity evaluations by the bioflocculanting assay of bacterial exopolysaccharides (EPS) produced by strain LZ-28 (pane A), and microalgae growth-promoting potential assay (pane B)

Genes coding enzymes for sulfur metabolism were present, *sox*A, *sox*B and *sox*C took part in the process of sulfur oxidation and electron transfer, *dmd*A, *dmd*B, *dmd*C and *dmd*D genes involved in DMSP (Dimethylsulfoniopropionate) demethylation, *dox*A encoding for hiosulfate: quinone oxidoreductase (TQO) subunits and *sir* encoding for sulfite reductase (SIR). For nitrogen metabolism, 10 genomes of *M. alba* strains also harbored genes encoding for dissimilatory nitrate reductase (*nar*G, *nar*H, *nar*I, *nar*J, *nar*K), and *ure*A, *ure*B, *ure*C, *ure*D, *ure*E, *ure*F, *ure*G encoding for urease showing the urea assimilation by the *M. alba* strains. The DMSP and amino acids provided by phycosphere microbiota will benefit the algal host’s nutrition acquirement during algae-bacteria interactions [34–37]. Moreover, stress response systems to defense oxidative stress were found in 10 genomes of *M. alba* strains. Reactive oxygen species (ROS) were released by algae in coping with different stressors during metabolism [38], and *M. alba* strains might be capable of coping with this oxidative stress in the algae-bacteria symbiotic system for survival as they harbored superoxide dismutase (SOD), catalases (CAT) and glutathione peroxidases (GPX), which played a role in scavenging oxygen free radicals and protecting cells from oxidative damage [39].

Quorum sensing (QS) signals were the means of communication among individuals living in a microbial community [40]. The genome analysis of strain LZ-28 predicted one autoinducer synthases (LuxI) and three LuxR type genes for AHL-controlled transcriptional regulators, along with two autoinducer 2 sensor kinase/phosphatase (*Lux*Q), three autoinducer 2 import system permease protein (*Lsr*D). *Lux*R were identified in *M. alba* genomes, which were related to AHL-mediated *Lux*R/I information system. QS signal molecules were thought to regulate the formation of biofilm, benifiting the attachment of organic particles and the successful colonization of bacteria. And the DMSP exchange between phycosphere microbiota and their dinoflagellates host that utilizes QS system [41–43].

## Discussion

The general features shown in Table 1 clearly indicated that the new isolate from phycosphere microbiota of toxic *Alexandrium catenella* LZT09, strain LZ-28 were similar to strains JLT354-W^T^, L6M1-5, JL351, F15 and KD53. For example, C_18 : 0_ and C_18 : 1_ *ω*7c 11-methyl were the predominant polar lipids of all strains of *Mameliella alba*. In addition, bacterial 16S rRNA gene similarity is also a direct evidence as golden standard for the taxonomy of bacterial species. Strain LZ-28 shared over 99.40% of 16S similarity with five representative *Mameliella alba* strains, together with additional evidences obtained by phylogenetic tree both confirming their close taxonomic relationship. Consequently, using the available genomes, three key phylogenomic parameters, including ANI, AAI and dDDH comparison were then performed. Species delimitation borders of ANI, AAI and dDDH analyses are generally set at 95-96%, 96% and 70%, respectively. Based on the genome evolution, relatedness of ANI, AAI and dDDH values, strain LZ-28 is a similar genomic species with *Mameliella alba* with the values ranges of 97%-100%, 97%-100% and 83%-100% compared with *Mameliella alba* strains, respectively. Taking the morphological and biochemical characterization and phylogenetic analysis together, strain LZ-28 could be considered as a new member of *Mameliella alba*.

For bacterial isolation, four strains including LZ-28, KD53, UMTAT08 and EP20 were all isolated from marine microalgaes, including three marine dinoflagellates, toxic *Alexandrium catenella* LZT09 in this study, *Alexandrium tamiyavanichii* AcMS01 [45] and *Symbiodinium* sp. [27], and also a marine diatom *Phaeodactylum tricornutum* [12], respectively. This interesting finding may indicate they were involved in potential symbiotic associations between *Mameliella* spp. with various host microalgaes [1–3,30]. Additionally, we found *Mameliella* spp. was among the top 10 abundant genus distributed in six HAB dinoflagellates obtained by metagenomics (Fig. S1) [7–10,18,19]. The equilibrium state of the symbiotic system was dynamically maintained by the regulation of numerous functional genes [23]. In this study, several BGCs responsible for synthesis of bacterial secondary metabolites including carotenoid, ectoine, type 1 polyketide synthases (PKS), bacteriocin and vitamin B_12_, respectively, were identified among the 10 strains of *M. alba* (Fig. 7). In a previous study, terpene was confirmed to play an important role in the symbiosis between the microbiota and its host dinoflagellate [24]. Terpene was an essential pigment for photosynthesis since it was closely related to the synthesis of bacterial carotenoids [25]. Similarly, bacteriocins have been considered to protect the algal host against pathogenic bacteria by developing resistance to specific pathogens [44].

Carbohydrates and energy are produced in photosynthesis, and thus well benefit for the bacterial adaption to harsh environment when lacking of sufficient nutrition for bacteria. In addition, micronutrients not only were delivered from phytoplankton to bacteria, but also usually the vice versa. Examples are essential organic compounds, such as thiamine (vitamin B_1_) and cobalamin (vitamin B_12_) acting as key compounds of enzyme cofactors, but most HAB algal species are vitamin B_1_ and B_12_ auxotrophs [32]. Vitamin B_12_ supplied by strain LZ-28 could maintain a stable growth state during algae-bacterial interactions since there is a specific demand for vitamin B_12_ on phytoplankton, and vitamins can have important ecological relevance for HAB [33].

The role of vitamin B_12_ in algal metabolism is primarily as a cofactor for vitamin B_12_-dependent synthase for methionine (MET) which is a necessary key precursor substance for initiating the PSTs biosynthesis [46]. However, vitamin B12, which is only synthesized *de novo* by some bacteria and archaea, is derived from uroporphyrinogen III, a precursor in the synthesis of heme, siroheme, cobamides, chlorophylls and the methanogenic F430 [51]. The genome of strain LZ-28 harbors all the genes required for *de novo* B_12_ synthesis. It’s similar to other genomic sequenced members of *Mameliella alba*.

## Conclusions

Combined polyphasic taxonomic analysis conducted in this study revealed that strain LZ-28 which was isolated from highly-toxic marine phycosphere is a new member of *Mameliella alba*. Comparative genomic study of the genus of *Mameliella* indicates the genomic potential for the cross-kingdom associations between *Mameliella* strains and their algal hosts. Moreover, stress response systems to defense oxidative stress and quorum sensing (QS) signals jointly maintain the balance and stability of the symbiotic environment between bacteria and their algal hosts. Furthering deeply digging the multiomics information obtained by *M. alba* strains and the algae host with interactive co-culture circumstance will provide hopeful insights on detailed mechanisms governing these dynamic algae-bacterial interactions occurred within the unique marine phycosphere niches.

## Supporting information

**Fig. S1.** Microbial abundance distribution at family level (left) and the top ten genus within the family *Rhodobacteraceae* (right) in six HAB dinoflagellates *Alexandrium* spp. obtained by metagenomics analysis. *Alexandrium tamarense* AT-11, *Alexandrium tamarense* AT 260, *Alexandrium minutum* AMKS5, *Alexandrium minutum* AMTK4, *Alexandrium catenella* ACHQ and *Alexandrium catenella* ACHK.

**Fig. S2.** Phylogenetic tree based on 16S rRNA showing the relationship between different strains belonging to *Mameliella* genus and type strains within the *Rhodobacteraceae* family by using maximum-likelihood (ML) algorithm. Bootstrap values (≥ 50%) based on 1000 replications are shown at the branch points. *Bar*, 0.01 substitutions per site. *Rhodococcus imtechensis* RKJ300^T^ was used as an outgroup.

**Fig. S3.** Phylogenetic tree based on 16S rRNA showing the relationship between different strains belonging to *Mameliella* genus and type strains within the *Rhodobacteraceae* family by using maximum-parsimony (MP) algorithm. Bootstrap values (≥ 50%) based on 1000 replications are shown at the branch points. *Bar*, 0.01 substitutions per site. *Rhodococcus imtechensis* RKJ300^T^ was used as an outgroup.

**Fig. S5.** Pan phylogenetic tree based on the pan-genome of 10 *Mameliella* strains generated by BPGA pipeline. The scale bar indicates time period in millions of years (MYA).

**Fig. S6.** Core phylogenetic tree based on the 4,472 core genes of 10 strains of *Mameliella alba* generated by BPGA pipeline. The scale bar indicates time period in millions of years (MYA).

**Table S1**. Detailed information about genome sequences of 10 strains of *Mameliella alba*.

## Abbreviations

HAB: Harmful algal blooms
EPS: Exopolysaccharides
ANI: Average nucleotide identity
AAI: Average amino acid identity
dDDH: digital DNA-DNA hybridization
QS: Quorum sensing
GTAs: Gene transfer agents
ABI: Algae-bacteria interactions
PST: Paralytic shellfish poisoning toxins
PMP: Phycosphere Microbiome Project
PM: Phycosphere microbiota
DMSP: Dimethylsulfoniopropionate
MA: Marine agar
MGPB: Microalgae growth-promoting bacterium
PHB: Poly-β-hydroxybutyrate
PG: Phosphatidylglycerol
PE: Phosphatidylethanolamine
PC: Phosphatidylcholine
GL: Glycolipid
PL: Phospholipid
NJ: Neighbour joining
ML: Maximum likelihood
MP: Maximum parsimony
NCBI: National center for biotechnology information
UBCG: Up-to-date bacterial core gene
HGT: Horizontal gene transfer
BGCs: Biosynthetic gene clusters
PKS: Polyketide synthases
AHL: N-acylhomoserine lactone
PHA: Polyhydroxyalkanoate
DMSP: Dimethylsulfoniopropionate
TQO: Thiosulfate, quinone oxidoreductase
SIR: Sulfite reductase
ROS: Reactive oxygen species
SOD: Superoxide dismutase
CAT: Catalases
GPX: Glutathione peroxidases

## Author Contributions

Conceptualization: Qiao Yang.

Formal analysis: Wen-zhuo Zhu, Fei-fei Xu.

Funding acquisition: Xiao-ling Zhang.

Methodology: Fei-fei Xu, Yun Ye.

Supervision: Qiao Yang, Xiao-ling Zhang.

Writing - original draft: Wen-zhuo Zhu, Qiao Yang, Xiao-ling Zhang.

Writing - review & editing: Qiao Yang, Xiao-ling Zhang.

